# Fructose-1,6-bisphosphatase (FBPase) fine-tunes heterotrophic growth in cyanobacteria

**DOI:** 10.1101/2025.02.14.638281

**Authors:** Frauke Caliebe, Ravi Shankar Ojha, Marco Gruber, Marko Boehm, Lu Shen, Christopher Bräsen, Jacky L. Snoep, Karl Forchhammer, Martin Hagemann, Bettina Siebers, Kirstin Gutekunst

## Abstract

Cyanobacteria switch their carbon metabolism between photoautotrophy and heterotrophy during diurnal cycles. In cyanobacteria, the classical glycolytic control point is characterized by two catabolic phosphofructokinases (PFKs) and a bifunctional anabolic fructose-1,6-biphosphatase/sedoheptulose-1,7-biphosphatase (F/SBPase; *slr2094*) catalyzing two key reactions in the Calvin-Benson-Bassham (CBB) cycle. In addition, *Synechocystis* possesses a fructose-1,6-bisphosphatase (FBPase; *slr0952*) with yet unknown physiological function and biochemical properties. Hence, our aim was to investigate the FBPase and the interplay of the four enzymes in photoautotrophic and heterotrophic carbon metabolism.

We discovered that FBPase is specific for FBP, showing no SBPase activity, and unlike F/SBPase does not exhibit any biochemical regulatory properties. Growth studies with deletion mutants revealed that FBPase and PFKs play a major role under heterotrophic conditions. In contrast to F/SBPase, FBPase is not involved in the CBB cycle, but instead fine-tunes heterotrophic growth. Transaldolase cannot replace the function of SBPase in the CBB cycle.

In conclusion, the classical Embden-Meyerhoff-Parnass pathway control point, which is known to be mediated by the antagonistic enzyme pair PFK and FBPase in heterotrophic bacteria and eukaryotes, is also present in *Synechocystis*. We found redox-insensitive FBPases from plant chloroplasts to be closely related to *Synechocystis* FBPase, indicating that they might serve a similar function.

**Highlight:** *Synechocystis* fructose-1,6-bisphosphatase (*slr0952*) is unlike fructose-1,6-biphosphatase/sedoheptulose-1,7-biphosphatase (*slr2094*) monofunctional, not redox-regulated, and supports heterotrophy in darkness, despite catalyzing an anabolic reaction. Thus, presumably Slr2094 alone drives two Calvin-Benson-Bassham cycle key reactions.

## Introduction

Cyanobacteria are the only prokaryotes performing oxygenic photosynthesis and are widely recognized as ancestors of this process in the plant kingdom. During the day, cyanobacteria photosynthesize, fixing CO_2_ to generate carbohydrates via the Calvin-Benson-Bassham (CBB) cycle, also known as the reductive pentose phosphate (RPP) cycle, and store excess fixed carbon intracellularly as glycogen. In darkness, the direction of carbohydrate metabolism is reversed. In the heterotrophic mode, glycogen or extracellular glucose are degraded through various catabolic pathways, primarily the oxidative pentose phosphate (OPP) pathway and glycolysis via the catabolic Embden-Meyerhoff-Parnass (EMP) pathway. Besides having an either anabolic or catabolic carbohydrate metabolism, cyanobacteria also enter conditions under which mixed forms are required. Especially in transition stages between light and darkness or in the presence of external carbohydrates in the light, glycolytic pathways can be shortened to glycolytic shunts (PGI (phosphoglucose isomerase) and OPP shunt) that feed carbohydrates into the CBB cycle for replenishment to enhance CO_2_ fixation and growth (Makowka et al., 2020; Schulze et al., 2022). Similarly, in plants carbohydrate reserves from the vacuole and the cytosol are converted into ribulose-5-phosphate (Ru5P) via the so-called cytosolic glucose-6-phosphate (G6P) shunt, which comprises the first two enzymes of the glycolytic OPP pathway. Ru5P is then transported into the chloroplasts and further into the CBB cycle. Under stress conditions replenishment of the CBB cycle also occurs via a chloroplastidic G6P (OPP) shunt in plants (Sharkey, 2024; Sharkey & Weise, 2015; Xu et al., 2022).

The anabolic CBB cycle and the catabolic OPP and EMP pathways share several enzymes and reactions, enabling the opportunistic reversal of carbon flow based on environmental conditions such as light intensities, CO_2_ levels, and metabolic demands. While some non-rate-limiting enzymes function bidirectionally under physiological conditions, key enzymes catalyzing irreversible reactions are often tightly regulated to ensure precise control and fine-tuning of metabolic fluxes. The classical key control point for regulating the anabolic and catabolic directions of carbon flow, i.e. glycolysis and gluconeogenesis, is catalyzed by the antagonistic unidirectional enzyme couple phosphofructokinase (PFK) and fructose-1,6-bisphosphatase (FBPase) in bacteria and eukaryotes. PFK, recognized as key glycolytic enzyme, catalyzes the phosphorylation of fructose-6-phosphate (F6P) to fructose-1,6-bisphosphate (FBP) in the catabolic direction, and FBPase the reversed anabolic reaction, the dephosphorylation of FBP to F6P. Another important regulatory control point unique to photoautotrophs is the dephosphorylation of seduheptulose-1,7-bisphosphate (SBP) to seduheptulose-7-phosphate (S7P), catalyzed by SBPase (Fig. 1).

**Figure 1:**
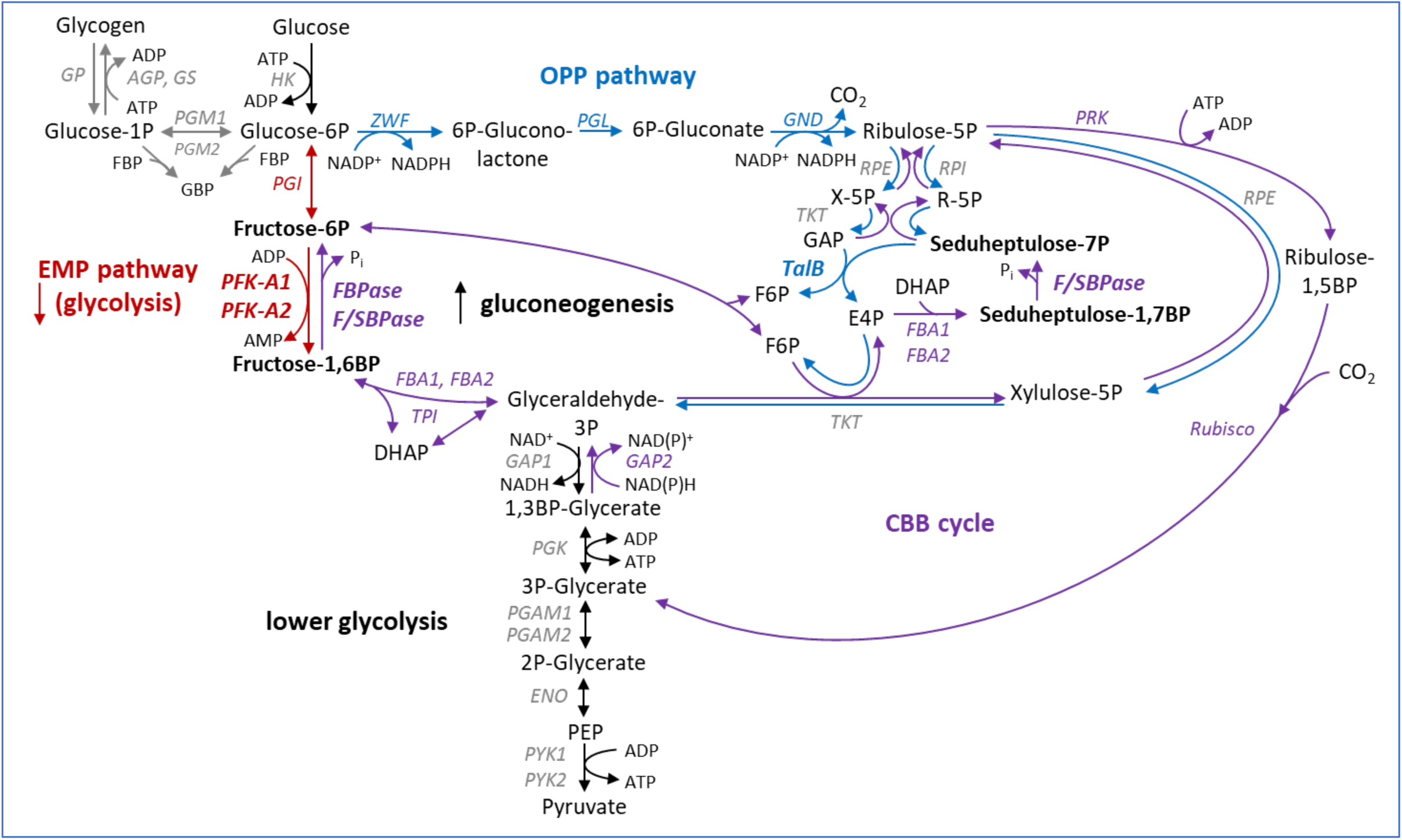
Reactions of the central carbohydrate metabolism in *Synechocystis*. Reactions of the CBB cycle are colored in purple, those of the OPP pathway in blue, and those of the EMP pathway in red. Many reactions of the CBB cycle overlap with the OPP or EMP pathway. (FBP = fructose-1,6-bisphosphate; F6P = fructose-6-phosphate; GBP = glucose-1,6-bisphosphate; X-5P = xylulose-5-phosphate; R-5P = ribose-5-phosphate; GAP = glyceraldehyde-3-phosphate; DHAP = dihydroxyacetone phosphate; PFK = phosphofructokinase; GP = glycogen phosphorylase; AGP = ADP-glucose pyrophosphorylase; GS = glycogen synthase; PGM = phosphoglucomutase; HK = hexokinase; ZWF = glucose-6-phosphate dehydrogenase; PGL = 6-phosphogluconolactonase; GND = 6-phosphogluconate dehydrogenase; RPE = ribulose-5-phosphate-3-epimerase; RPI = ribose-5-phosphate isomerase; TKT = transketolase; TalB = transaldolase; PGI = glucose-6-phosphate isomerase; FBPase = fructose-1,6-bisphosphatase; F/SBPase = fructose-1,6-biphosphatase/sedoheptulose-1,7-biphosphatase; PRK = phosphoribulokinase; FBA = fructose-bisphosphate aldolase; PYK = pyruvate kinase; ENO = enolase; PGAM = phosphoglycerate mutase; PGK = phosphoglycerate kinase; GAP = glycerinealdehyde-3-phosphate dehydrogenase; TPI = triosephosphate isomerase)

The latter reaction of SBPase is unique to the CBB cycle and has a strong control on the flux through the cycle (De Porcellinis et al., 2018). This notion is supported by findings that overexpression of SBPase, e.g. using a bifunctional cyanobacterial enzyme, increased the CO_2_ fixation and growth of plants (Miyagawa et al., 2001). No enzyme is known in cyanobacteria or plants that catalyzes the corresponding reverse reaction (the phosphorylation of S7P), whereas some Clostridia possess a PP_i_-PFK which is able to convert S7P to SBP (Koendjbiharie et al., 2020). However, transaldolase (TalB; *slr1793*), which is mainly attributed to the OPP pathway, is able to bypass the SBPase reaction in the catabolic direction, by converting glyceraldehyde-3-phosphate (GAP) and S7P to F6P and erythrose-4-phosphate (E4P) (Fig. 1). However, this reaction operates reversibly and is close to equilibrium. It is not well established if transaldolase is restricted to the OPP pathway or might in addition be involved in carbon fixation via the CBB cycle (Fridlyand & Scheibe, 1999). So-called transaldolase variants of the CBB cycle that lack SBPase and use transaldolase instead were reported in autotrophic bacteria as *Thermosulfobium acidiphilum* (Frolov et al., 2019). A similar pathway is likely used by *E. coli* strains that have been genetically engineered to grow autotrophically (Antonovsky et al., 2016).

Unlike plants, the cyanobacterial cell has no defined compartments. Hence all anabolic and catabolic routes of carbon metabolism occur in the cytoplasm, which increases the need of regulatory mechanisms to avoid futile cycles (reviewed in Lucius & Hagemann, 2024). Although two PFK isoenzymes, an FBPase (*slr0952*), and a bifunctional F/SBPase (*slr2094*) have been identified in *Synechocystis* sp. PCC 6803 (hereafter *Synechocystis*), this glycolytic control point was regarded to be absent in the organism (Knowles & Plaxton, 2003). However, our recent characterization of the *Synechocystis* PFK-A family isoenzymes, PFK-A1 (*sll1196*) and PFK-A2 (*sll0745*), revealed that both specifically utilize ADP as co-substrate instead of ATP and exhibit distinct allosteric regulation (Shen et al., 2024). PFK-A1 is inhibited by 3-phosphoglycerate (3PG), a product of the CBB cycle, while PFK-A2 is inhibited by ATP generated during the photosynthetic light reaction. Physiological experiments with *Synechocystis* showed diminished levels of polyhydroxybutyrate (PHB) production in the single mutants Δ*p[-A1* and Δ*p[-A2*, and the double mutant Δ*p[-A1*Δ*p[-A2* (called Δ*p[* from here on) under nitrogen starvation (Koch et al., 2019). In other experiments, the Δ*p[* mutant showed growth similar to that of WT under photoautotrophic and photomixotrophic conditions including different CO_2_ levels, but reduced growth under heterotrophic conditions (Chen et al., 2016; Lucius et al., 2021; Makowka et al., 2020). Growth data for the single Δ*p[-A1* and Δ*p[-A2* mutants are still lacking.

In contrast to plants, which use separate enzymes for dephosphorylating FBP and SBP during the CBB cycle within chloroplasts, *Synechocystis* employs a bifunctional F/SBPase. During the day, this enzyme is photoactivated by reduced thioredoxin, which breaks disulfide bonds between conserved cysteines in the enzyme. Furthermore, the enzyme is inhibited by AMP (K_i(FBP)_ 34 µM), ensuring activity in the light at high energy charge of the cell and minimal activity during darkness (Feng et al., 2014; Mallén-Ponce et al., 2021; Sporre et al., 2023; Tamoi et al., 1998). A similar photoactivation mechanism by reduction of cysteine-disulfide bonds is described for plant chloroplast cpFBPase and cpSBPase, but those are insensitive to AMP (Buchanan, 2016; Ladror et al., 1990; Nishizawa & Buchanan, 1981). Deletion of *slr2094* encoding F/SBPase in *Synechocystis* abolishes photoautotrophic growth, which could only be restored if both plant enzymes, the cpFBPase and cpSBPase were expressed in the mutant simultaneously (García-Cañas et al., 2022; Yan & Xu, 2008).

Notably, *Synechocystis* encodes an additional, distinct fructose-1,6-bisphosphatase (*fbpase*; *slr0952*). However, unlike F/SBPase, the function of this FBPase protein in *Synechocystis* remains enigmatic, as the enzyme has not been characterized, and no growth conditions have yet been identified under which FBPase plays a role. Deletion mutants of *slr0952* (Δ*fbpase*) consistently exhibited growth comparable to that of the wild type (WT) (García-Cañas et al., 2022). Furthermore, the expression level of *slr0952* (*fbpase*) is significantly lower than that of *slr2094* (*f/sbpase*) (Jackson et al., 2022). This led to the conclusion that FBPase *slr0952* is of minor importance.

Interestingly, plants possess a cytosolic cyFBPase in addition to the chloroplastic cpFBPase. This cytosolic enzyme plays a critical role by catalyzing the first irreversible step in sucrose synthesis, which is tightly regulated. Plant cyFBPases lack the conserved cysteine residues, and thus do not undergo photoactivation. However, they are inhibited by the carbon regulatory compound fructose-2,6-bisphosphate and AMP, similar to gluconeogenetic FBPases from mammals and yeast (Khayat et al., 1993; Ladror et al., 1990; Stitt et al., 1985; Zhou & Cheng, 2004). In some plants, a second chloroplastic cpFBPase II has been identified, which also lacks the conserved cysteine residues required for photoactivation. However, its physiological role remains unclear (Li et al., 2020; Serrato et al., 2009).

The aim of this study is to gain a deeper understanding of the critical role played by the antagonistic enzyme pairs PFK-A1, PFK-A2 and FBPase, F/SBPase and TalB in regulating the switch between catabolic and anabolic carbon metabolism in cyanobacteria. Specifically, we seek to clarify the enzymatic regulation and function of FBPase, as well as its physiological role through mutant studies.

## Materials and Methods

### Gene cloning and protein overexpression

The FBPase (*slr0952*) and F/SBPase encoding genes (*slr2094*) were amplified from *Synechocystis* sp. PCC 6803 genomic DNA, using the primer sets *slr0952*_forward_NdeI_5’ & *slr0952*_reverse_BamHI_5’ and *slr2094*_forward_XhoI_5’ & *slr2094*_reverse_BamHI_5’, respectively (for sequences, see Table S1). Subsequently, *slr0952* and *slr2094* were cloned into the expression vector pET15b with N-terminal 6xhistidine-tag (Novogene, Beijing, China). Successful cloning was confirmed by DNA sequencing (LGC genomics, Berlin, Germany). The expression plasmid (pET16b_bmmga3_16125) for fructose-1,6-bisphosphate aldolase from *Bacillus methanolicuss* (*Bm*FBA) used for synthesis of sedoheptulose-1,7-bisphosphate (SBP) was kindly provided by V. Wendisch (Universität Bielefeld) (Stolzenberger et al., 2013b). For expression, the respective plasmids were transformed into the *Escherichia coli* (*E. coli*) strain Rosetta (DE3, Stratagene) and overexpression was performed in 1 L terrific broth (TB) medium (yeast extract 22 g/L, tryptone 12 g/L and glycerol 4 mL/L) containing 100 µg/mL ampicillin and 30 µg/mL chloramphenicol. Cells were cultivated at 37°C with shaking at 180 rpm (Unitron, INFORS HT, Bottmingen, Switzerland) and protein expression was induced at an OD_600_ of 0.6-0.8 by addition of 1 mM isopropyl-β-D-thiogalactopyranoside (IPTG). After induction, the cultures were further incubated at 18°C and 180 rpm for 18-22 h. Cells were collected by centrifugation (15 min, 8630 x g, 4 °C) and stored at -70°C until use.

### Protein purification

Both the recombinant *Synechocystis* FBPase and F/SBPase, as well as *Bm*FBA were purified by immobilized metal ion affinity chromatography (IMAC) and size exclusion chromatography (SEC). For purification, frozen cells from FBPase, F/SBPase and *Bm*FBA expression were resuspended in 50 mM HEPES/NaOH (pH 7.8, 30°C), 300 mM NaCl (1 g cells (wet weight)/ 3 mL buffer). The cells were disrupted using sonicator (3 x 5 minutes, amplitude 50, cycle 0.5) (UP200S, Hielscher Ultrasonics, Brandenburg, Germany). Cell debris was removed by centrifugation (45 min, 21,130 x g, 4°C) and the histidine-tagged proteins were purified from the supernatant using nickel (tris(carboxymethyl)ethylenediamin) columns (Ni-TED) columns (Macherey-Nagel, Dueren, Germany) according to the manufacturers’ instructions. Elution fractions containing the recombinant protein were collected and concentrated using centrifugal concentrators (Vivaspin®20, Satorius Stedium Biotech, cut off size 30 kDa). Afterwards, the concentrated protein samples of either FBPase, F/SBPase or *Bm*FBA (6, 3.5, and 6 mg, respectively) were applied onto a SEC column (HiLoad 16/600 Superdex 200 prep grade, Amersham Biosciences) pre-equilibrated with 50 mM HEPES/NaOH (pH 7.8, 30°C), 300 mM NaCl. Protein fractions for FBPase, F/SBPase or *Bm*FBA were collected after SEC (5, 3 and 4.5 mg, respectively) and analyzed by activity measurements and SDS-PAGE. Proteins were stored at -70°C in the presence of 25% (v/v) glycerol. The protein concentration was determined using the Bradford assay (Zor & Selinger, 1996) with BSA (Merck, Darmstadt, Germany) as standard.

### Determination of the native molecular mass

After purification by IMAC, 1 mg of FBPase was loaded on to a SEC column (superose 6 Increase 10/300 GL column, Amersham Biosciences). As divalent metal requirement is reported for other FBPases (Donahue et al., 2000; Stolzenberger et al., 2013a) we observed an effect of MgCl_2_ on the oligomeric structure of FBPase. After 6 hours of incubation with 10 mM MgCl_2_ in 50 mM HEPES/NaOH (pH 7.8, 30°C), 300 mM NaCl, the enzyme separation revealed two peaks, representing either a dodecameric or tetrameric structure. After 48 hours incubation, only the tetrameric form was observed. Using the same SEC column, the calibration curve was generated with five proteins (aprotin (6.5 kDa), ovalbumin (43 kDa), aldolase (158 kDa), ferritin (440 kDa) and thyroglobulin (669 kDa) and blue dextran from the LMW and HMW gel filtration calibration kits (GE Healthcare), using the same buffer as for the purification of recombinant FBPase. The native molecular mass of FBPase was calculated using the generated calibration curve.

### *In vitro* FBPase activity measurements

The FBPase activity of FBPase and F/SBPase was determined using a continuous, coupled enzymatic assay to monitor the conversion of fructose-1,6-bisphosphate (FBP) to fructose-6-phosphate (F6P). This reaction was facilitated by two auxiliary enzymes: phosphoglucoisomerase (PGI, from *Saccharomyces cerevisiae;* Merck, Darmstadt, Germany), which converts the formed F6P to G6P, and glucose-6-phosphate dehydrogenase (G6PDH, rabbit muscle; from *Saccharomyces cerevisiae;* Merck, Darmstadt, Germany), which oxidizes G6P to 6-phosphogluconate (6PG) while reducing NADP^+^ to NADPH. The increase in NADPH was detected by measuring the absorbance at 340 nm in 96-well plates (BRANDplates®, BRAND, Wertheim, Germany). A NADPH calibration curve (0 – 0.7 mM NADPH) was used for quantification and measurements were conducted using a Tecan Infinite M200 plate reader (Tecan Group AG, Männedorf, Switzerland) at 30°C.

The assay mixture (200 µL total volume) contained 1 μg FBPase or 1.6 μg F/SBPase, 50 mM HEPES/NaOH (pH 7.8 at 30°C), 1 U PGI, 1 U G6PDH, 10 mM MgCl_2_, 5 mM NADP^+^ and 3 mM FBP. For F/SBPase 10 mM DTT was included in the assay. The reaction was started by adding the enzyme.

To determine the metal ion dependency, various divalent metal ions were tested at concentrations of 1 and 5 mM using the standard assay. For the best performing metal ions, the optimal concentration was determined, for MgCl_2_ within a range of 1 - 20 mM, for MnCl_2_ in a narrower range of 0.01 - 5 mM. The enzyme characterization was performed in the presence of 10 mM MgCl_2_. To calculate V_max_ and K_m_ values, the Michaelis-Menten equation was fit using the NonlinearModelFit function in Wolfram Mathematica v14.

Effector studies were performed using the standard assay in the presence of 1 mM fructose-1-phosphate (F1P), phosphoenolpyruvate (PEP), citrate, isocitrate, malate, 2-phosphoglycolate (2PG), ATP, ADP, AMP and DTT with a half-saturation concentration of FBP (0.2 mM). A more detailed characterization was performed for ATP, AMP and DTT, using 1-10 mM of the effectors. Additionally, for ATP, the experiment was repeated with 30 mM MgCl_2_.

All assays were performed in triplicate. Negative controls were conducted by omitting either enzyme or FBP. One unit (1 U) of enzyme activity is defined as 1 µmol of NADP^+^ being reduced to NADPH via the auxiliary enzyme G6PDH per minute.

### *In vitro* SBPase activity measurements

Since SBP was not commercially available, *Bm*FBA was employed for synthesis. *Bm*FBA catalyzes the reversible conversion of FBP to glyceraldehyde-3-phosphate (GAP) and dihydroxyacetone phosphate (DHAP), as well as the reversible condensation of erythrose-4-phosphate (E4P) and DHAP to SBP (Stolzenberger et al., 2013b). The activity of *Bm*FBA was confirmed in cleavage direction using FBP as the substrate. The reaction mixture (200 µL total volume) consisted of 2 μg *Bm*FBA, 50 mM HEPES/NaOH (pH 7.8 at 30 °C), 0.7 mM NADH, 5 mM FBP and 1 U glycerol-3-phosphate dehydrogenase (G3PDH, from rabbit muscle; Merck, Darmstadt, Germany). In the cleavage reaction, DHAP, one of the products, was converted to glycerol-3-phosphate (G3P) resulting in the oxidation of NADH to NAD^+^. The oxidation of NADH was monitored as a decrease in absorbance at 340 nm in 96-well plates (BRANDplates®, BRAND, Wertheim, Germany). A NADPH calibration curve (0 – 0.7 mM NADPH) was used for quantification, and measurements were conducted using a Tecan Infinite M200 plate reader (Tecan Group AG, Männedorf, Switzerland) at 30°. A specific activity of 4 Umg^-1^ was determined for *Bm*FBA using this assay.

To analyze the SBPase activity of FBPase, three different assays were performed. Each reaction contained 3 mM E4P, 3 mM DHAP and 50 mM HEPES/NaOH (pH 7.8 at 30 °C). The specific compositions of the reactions were as follows: reaction A included 10 μg *Bm*FBA; reaction B 10 μg *Bm*FBA, 10 mM MgCl_2_ and 5 μg FBPase; and reaction C 10 μg *Bm*FBA, 10 mM MgCl_2_, 10 mM DTT and 5 μg F/SBPase. The reactions were incubated at 30°C for 2 hours. After incubation each reaction was diluted 30-fold to achieve a final concentration of 0.1 mM of substrates/ products. From the diluted mixtures, 0, 10, 20, 30, 40 and 50 μl of the reaction were mixed with 30 μl of malachite green phosphate reagent (Merck, Phosphate calorimetric assay kit; Darmstadt, Germany) and H_2_O was added to a final volume of 200 μL. Absorbance at 650 nm in 96-well plates (BRANDplates®, BRAND, Wertheim, Germany) was measured using a Tecan Infinite M200 plate reader (Tecan Group AG, Männedorf, Switzerland).

### 31P NMR analysis of FBPase and F/SBPase

For qualitative analysis, the enzyme reactions of FBPase and F/SBPase were analyzed via ^31^P NMR spectroscopy. Control samples, including 3 mM FBP, 3 mM phosphate (PO_4_^3-^) or 3 mM F6P, were prepared in 50 mM HEPES/NaOH (pH 7.8, 30°C), 300 mM NaCl, and 10 mM MgCl_2_. The enzyme assays were performed in the same buffer with 3 mM FBP, and 5 µg FBPase or F/SBPase. For F/SBPase, the reaction mixture also included 10 mM DTT. All reaction mixtures were incubated at 30° C for 2 hours. The ^31^P NMR spectra was obtained using a Bruker Avance Neo 400 device, and data visualization was performed using TopSpin 3.7.0 software.

### Bioinformatic analysis

Structural models were obtained from the AlphaFold Server version 3.0 (Abramson et al., 2024). Structural analyses, comparative assessments, and visualizations were conducted using the UCSF ChimeraX software suite, developed by the Resource for Biocomputing, Visualization, and Informatics at the University of California, San Francisco (Pettersen et al., 2021). Amino acid sequences for sequence alignment were retrieved from the UniProt database, and alignments were generated using the EMBL-EBI Job Dispatcher framework for sequence analysis tools (Madeira et al., 2024). The BioEdit software suite was used for manual refinement and editing of sequence alignments. Protein alignments of selected FBPase and SBPase proteins from heterotrophic bacteria, cyanobacteria, green algae and streptophytes were used for phylogenetic analyses using the software package MEGA. The protein sequence of GlpX from *Corynebacterium glutamicum* served as outgroup. For GenBank accession numbers of the used sequences, refer to Table S2.

### Generation and verification of deletion mutants of *Synechocystis*

To generate deletion mutants, *Synechocystis* was transformed with constructs, in which a resistance cassette was fused to around 200 bp directly up- and downstream of the gene to be deleted. The fragments for these constructs were amplified by PCR and subsequently cloned into pBluescript vector by Gibson Assembly (Gibson et al., 2009); alternatively, a pUC vector already containing the up- and downstream regions was ordered from Genewiz (GENEWIZ Germany GmbH, Leipzig, Germany), and opened with EcoRV between the up- and downstream regions, and the resistance cassette amplified by PCR was inserted by Gibson Assembly.

For testing the segregation of the mutants, a PCR was performed (Supplemental Text S1, Figs. S1-5).

All mutants that were used are listed in Table S3, along with the primers that were used for their generation and for testing their segregation. Some of them had already been generated previously. All primer sequences are listed in Table S1.

### Growth conditions for *Synechocystis*

Strains were maintained on BG11 plates at 28°C under constant illumination (50-100 µmol/(m²s), mutants with antibiotics.

For growth experiments, *Synechocystis sp.* PCC 6803 WT and mutants were grown in 200 mL BG11 medium (pH 8) at 28°C in glass tubes and equally aerated with filter-sterilized air, as described previously (Makowka et al., 2020). To monitor growth, OD_750_ was measured every day, using a 96-well plate and a plate reader (Infinite M Nano+, Tecan Group AG, Männedorf, Switzerland). In each experiment, three biological replicates were used and independent experiments for every analyzed condition were conducted at least three times. Pre-cultures for growth experiments were grown autotrophically, first in baffled shake flasks for one day (in 50 mL BG11), then in glass tubes for further six days (diluted with 50 mL BG11). For mutants, antibiotics were used in the pre-cultures, but not in the growth experiments, unless noted otherwise. Cultures were illuminated from two sides with constant light of 50 µmol/(m²s); only heterotrophic cultures were kept in darkness, except for daily illumination of around 10 minutes. For photomixotrophic, photoheterotrophic and heterotrophic cultures, 10 mM glucose was added. For photoheterotrophic cultures, 40 µM 3-(3,4-dichlorophenyl)-1,1-dimethylurea (DCMU) was added additionally.

The following antibiotic concentrations were used: kanamycin 50 µg/mL; spectinomycin 20 µg/mL; gentamycin 10 µg/mL in agar plates, 2.5-10 µg/mL in liquid culture.

### Determination of FBPase activity in crude cell extracts of *Synechocystis*

The method for measuring FBPase activity in crude cell extracts of *Synechocystis* was based on (Yan & Xu, 2008), but some adaptations were made. Like in the *in vitro* measurements, the substrate of FBPase (fructose-1,6-bisphosphate) was supplied and phosphoglucoisomerase and glucose-6-phosphate dehydrogenase were used as auxiliary enzymes to convert its product (fructose-6-phosphate) to 6-phosphogluconate. The NADPH generated in the last step was quantified by measuring the absorbance at 340 nm.

Cells from 50 mL liquid culture were harvested by centrifugation (3,000 rpm, 10 min, 4°C), the pellet was washed in potassium phosphate buffer once (50 mM, pH 8.0; 6,000 g, 10 min, 4°C) and the pellet was resuspended in 2 mL extraction buffer (50 mM potassium phosphate pH 8.0, 2.5 mM DTT, 1 mM glutathione, 10% v/v sucrose). The suspension was distributed into 2 reaction tubes containing around 400 µL glass beads (0,17-0,18 mm) and vortexed (3 min, 4°C). Two centrifugation steps followed (maximum speed, 1 min, 4°C; then 12,000 rpm, 20 min, 4°C). The supernatant (crude extract) was diluted with extraction buffer to the same phycobilisome concentration for all samples, determined by measuring the absorbance at 650 nm. 20 µL crude extract were mixed with 160 µL enzyme mix (100 mM Tris-HCl pH 8.0, 10 mM MgCl_2_, 0.5 mM EDTA, 1.5 mM Na_2_NADP, 0.5 U/ml glucose-6-phosphate dehydrogenase, 1.5 U/ml phosphoglucoisomerase, 10% v/v glycerol). The baseline absorption at 340 nm was measured over 10 minutes in a 96-well plate with a plate reader (Infinite M Nano+, Tecan Group AG, Männedorf, Switzerland). 20 µL substrate mix were added (100 mM Tris-HCl pH 8.0, 10 mM MgCl_2_, 0.5 mM EDTA, 20 mM fructose-1,6-bisphosphate), and after shaking, the NADPH production was quantified by measuring the absorbance at 340 nm over 30 minutes. FBPase enzyme activity was quantified with a calibration curve of NADPH (0 – 0.16 µM).

For testing reducing conditions, the measurements were conducted as described, for oxidizing conditions, extraction buffer without DTT was used. We verified that the presence or absence of DTT did not influence the auxiliary enzymes and that extraction without DTT did indeed represent reducing conditions (Supplemental Text S2, Fig. S6). Furthermore, we observed that addition of DTT did not increase the activity after extracting under oxidizing conditions, which indicates that the reduction for activating F/SBPase requires a cellular mediator (Supplemental Text S2, Fig. S7).

To test for significant differences, a one-way analysis of variance (ANOVA) was performed for the two influence factors strain and extraction method, for the whole dataset and also stratified on the other influence factor. If the factor strain was significant, posthoc comparisons were performed according to the Tukey methods which means that these are adjusted for multiple testing. The significance level for all tests was 0.05 and all tests were two-sided. All analyses were performed with the statistics software R, version 4.4.0 (R Core Team, 2024. R: A Language and Environment for Statistical Computing. R Foundation for Statistical Computing, Vienna, Austria. https://www.R-project.org/).

## Results

### Overexpression and purification of the recombinant FBPase

For the biochemical characterization of the FBPase from *Synechocystis*, the encoding gene (*slr0952, fbpase*) was cloned in the pET expression system with an N-terminal His-tag. The recombinant enzyme was expressed in *E. coli* Rosetta (DE3) cells and subsequently purified (Fig. S8). Approximately 6 mg of pure FBPase was obtained from 2.6 g cells (wet weight). The molecular mass of FBPase under denaturing conditions in SDS-PAGE was approximately 40 kDa, which aligns well with the calculated molecular mass of 39 kDa. The native molecular mass after preincubation with 10 mM MgCl_2_, determined by size-exclusion chromatography (SEC), showed a single peak at 163 kDa, suggesting a homotetrameric structure for FBPase (Fig. S9). This observation is consistent with previous studies reporting that class I FBPases typically exhibit a homotetrameric structure (Hines et al., 2006).

### Biochemical characterization of FBPase

Like most known FBPases (Brown et al., 2009; Stieglitz et al., 2003), *Synechocystis* FBPase is a metal-dependent enzyme that requires divalent metal ions for activity. In the absence of divalent metal ions, the enzyme showed no activity with FBP as the substrate (Fig. S10A). Significant FBPase activity was observed only in the presence of Mg^2+^. Mn^2+^ and Co^2+^ could partially activate the enzyme, though to a lesser extent. This metal response differs from previously characterized bifunctional F/SBPases, such as the one from *Synechocystis*, which can utilize both Mg^2+^ and lower concentrations of Mn^2+^ to shift from an inactive homodimeric form to an active tetrameric form (Feng et al., 2014; Hines et al., 2006). In contrast, *Synechocystis* FBPase specifically requires Mg^2+^. The inclusion of 10 mM Mg^2+^ in the reaction buffer resulted in the highest FBPase activity, reaching approximately 12 U mg^-1^ protein (Fig. S10B). To determine kinetic parameters, FBPase was incubated with varying FBP concentration. The enzyme follows a classical Michaelis-Menten kinetics with a V_max_ of 12.7±0.19 U mg^-1^ protein and a K_m_ of 0.18±0.01mM (Fig. 2). In comparison to F/SBPase, FBPase has a 2-fold lower affinity towards FBP (K_m_) and catalyzes its dephosphorylation with a slightly lower turnover number (K_cat_), resulting in lower catalytic efficiency (K_cat_/K_m_) values (Table 1).

**Figure 2:**
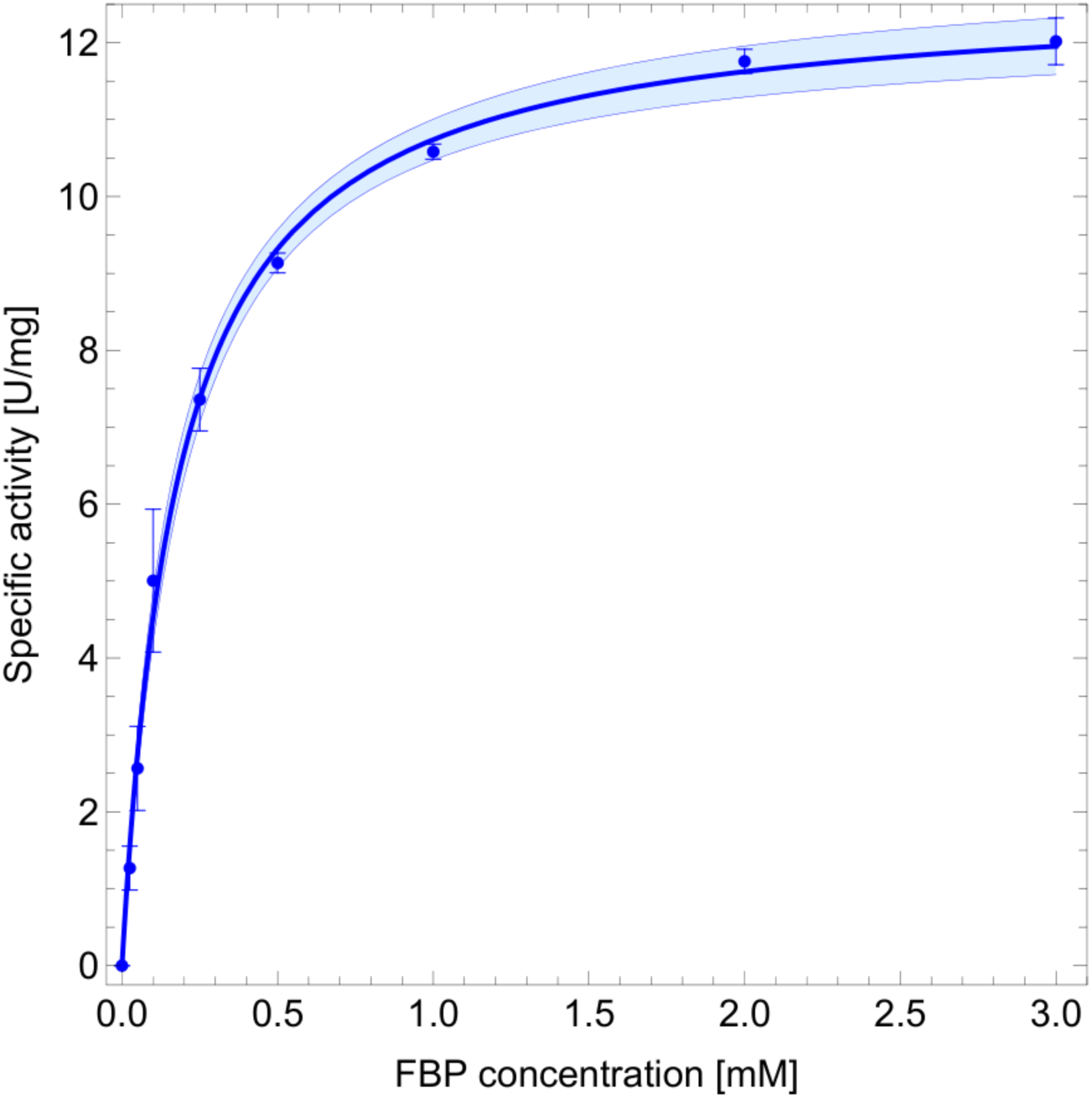
Enzymatic characterization of FBPase. The enzymatic activity of FBPase was assessed with varying concentrations of FBP (0-3 mM) in the presence of 10 mM MgCl_2_. Data are presented as means ± standard deviations from three independent measurements (technical replicates) of two biological replicates (n = 6). The shaded region shows the 95% confidence interval. To determine V_max_ (12.7 U mg^-1^) and K_m_ (0.18 mM) values, the Michaelis-Menten equation was fit using the NonlinearModelFit function in Wolfram Mathematica v14.2.

**Table 1:** Kinetic parameters of the *Synechocystis* F/SBPase and FBPase.

| Parameters | F/SBPase (Feng et. al. 2014) | FBPase (this work) |
| --- | --- | --- |
| K<sub>cat</sub> (FBP) | 10.5 sec<sup>-1</sup> | 8.4 sec<sup>-1</sup> |
| K<sub>cat</sub> (SBP) | 4.2 sec<sup>-1</sup> | NA |
| K<sub>cat</sub>/K<sub>m</sub> (FBP) | 131.25 sec<sup>-1</sup>mM<sup>-1</sup> | 46.7 sec<sup>-1</sup> mM<sup>-1</sup> |
| V<sub>max</sub> (FBP) | 16.1 U mg<sup>-1</sup> * | 12.7 U mg<sup>-1</sup> |
| K<sub>m</sub> (FBP) | 0.08 mM | 0.18 mM |
| K<sub>m</sub> (SBP) | 0.24 mM | NA |
| Activators | DTT | None |
| Inhibitors | AMP (K<sub>i</sub>(FBP) = 34 μM) | None |

NA = not active
\* = Determined by authors using the given K<sub>cat</sub> value in Feng et. al. 2014

Previous studies on class I FBPases have demonstrated that several metabolic intermediates such as fructose-1-phosphate, malate, citrate, isocitrate, and AMP can inhibit the enzyme (Babul & Guixé, 1983; Van Praag, 1997; Donahue et al., 2000; Wolf et al., 2018). The intriguing regulation of the *Synechocystis* F/SBPase isoenzyme, which is activated by DTT and inhibited by AMP, has revealed its relevance under light conditions (Feng et al., 2014), and serves as a foundation for investigating the biochemical regulation of *Synechocystis* FBPase as well. In this study, we tested several effectors under half saturating conditions for FBP. None of the tested effectors caused a significant effect, except for a slight reduction in activity by ATP, DTT and AMP (93, 91, and 94% residual activity, respectively, in the presence of 1 mM of these effectors) (Fig. 3). To further explore these inhibitory effects, we varied the concentration of ATP, DTT and AMP under half saturating FBP conditions. ATP was the only effector that caused significant inhibition with a K_i_ of 5.3 mM (Fig. S11). However, further studies with increased Mg^2+^ concentrations (30 mM) revealed that the inhibitory effect of ATP was due to competition for Mg^2+^ required for enzyme activity. Hence, ATP inhibition of the *Synechocystis* FBPase is probably not relevant under *in vivo* conditions. In summary, none of the tested effectors had a regulatory effect on FBPase. Notably, AMP and DTT, which are known to affect the activity of *Synechocystis* F/SBPase, did not impact the activity of FBPase, even at concentrations up to 10 mM.

**Figure 3:**
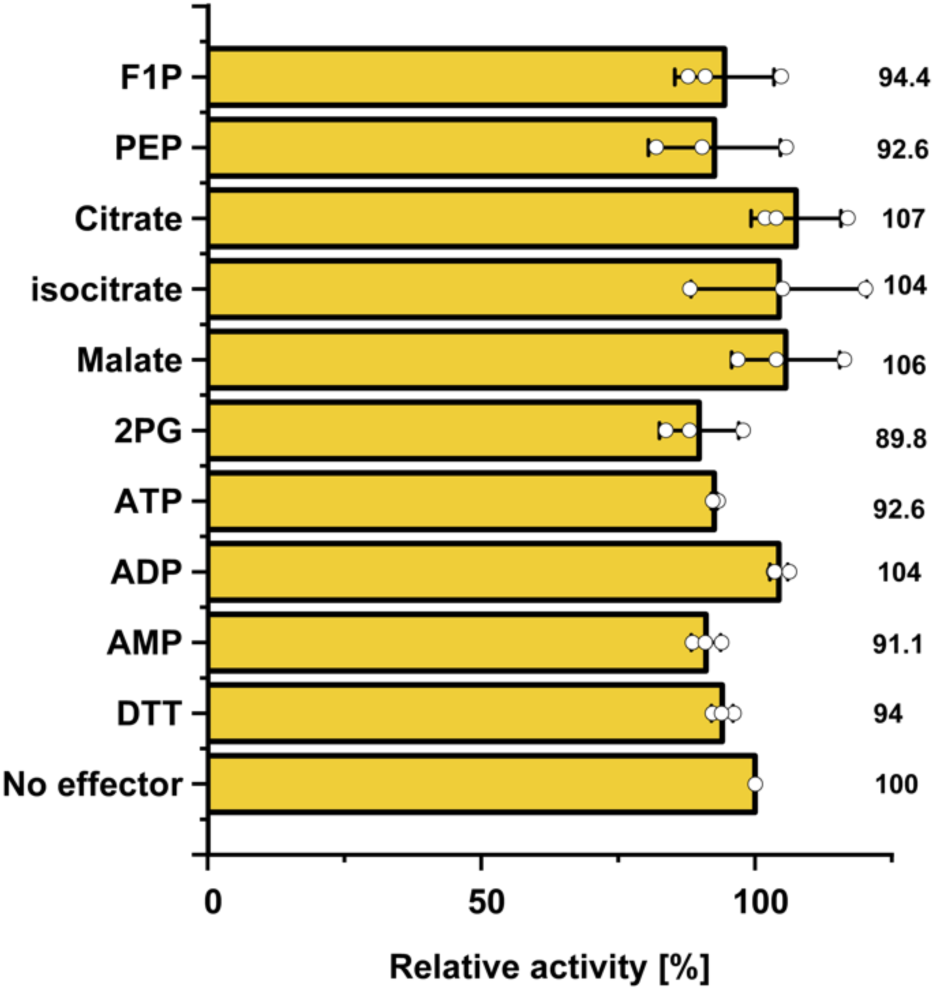
Influence of different metabolites and effectors on FBPase activity. The effects of various metabolites and effectors (1 mM) on FBPase activity was assessed under subsaturating (0.2 mM FBP) conditions. Relative activity (%) is expressed as a percentage of the control activity measured in the absence of effector (100%). The specific activities of the control under subsaturating conditions was 6 U mg^-1^. Data represent the means ± standard deviations from three technical replicates (n = 3).

### Monofunctional FBPase lacks SBPase activity

To test if the *Synechocystis* FBPase is mono- or bifunctional, we compared its possible SBPase activity with the bifunctional F/SBPase as positive control. First, the FBPase activity of recombinant F/SBPase was confirmed in the presence of 10 mM Mg^2+^ and 10 mM DTT. Consistent with previous reports (Feng et al., 2014), its activity was almost completely inhibited by 50 µM AMP (Fig. S12). Additionally, ^31^P NMR analysis clearly verified that both, FBPase and F/SBPase are able to irreversibly cleave FBP to F6P and PO_4_^3-^ (Fig. S13).

To test the SBPase activity of the enzymes, the substrate SBP was produced by recombinant fructose-1,6-bisphosphate aldolase from *Bacillus methanolicus* (*Bm*FBA), as SBP was not commercially available. *Bm*FBA catalyzes the reversible condensation of E4P and DHAP to SBP (Stolzenberger et al., 2013b). Only in the presence of the *Synechocystis* F/SBPase we observed release of PO_4_^3-^ using a malachite green assay (Fig. 4A), whereas the FBPase did not cleave SBP. Additionally, we verified that DHAP consumption, indicative of SBPase activity, only occurred in the presence of both *Bm*FBA and F/SBPase, but neither with F/SBPase alone nor with *Bm*FBA and FBPase from *Synechocystis* (Fig. 4B). In contrast, both enzymes did not accept glucose-1,6-bisphosphate as substrate, and it did also not act as effector for FBPase or F/SBPase activities (Fig. S14).

**Figure 4:**
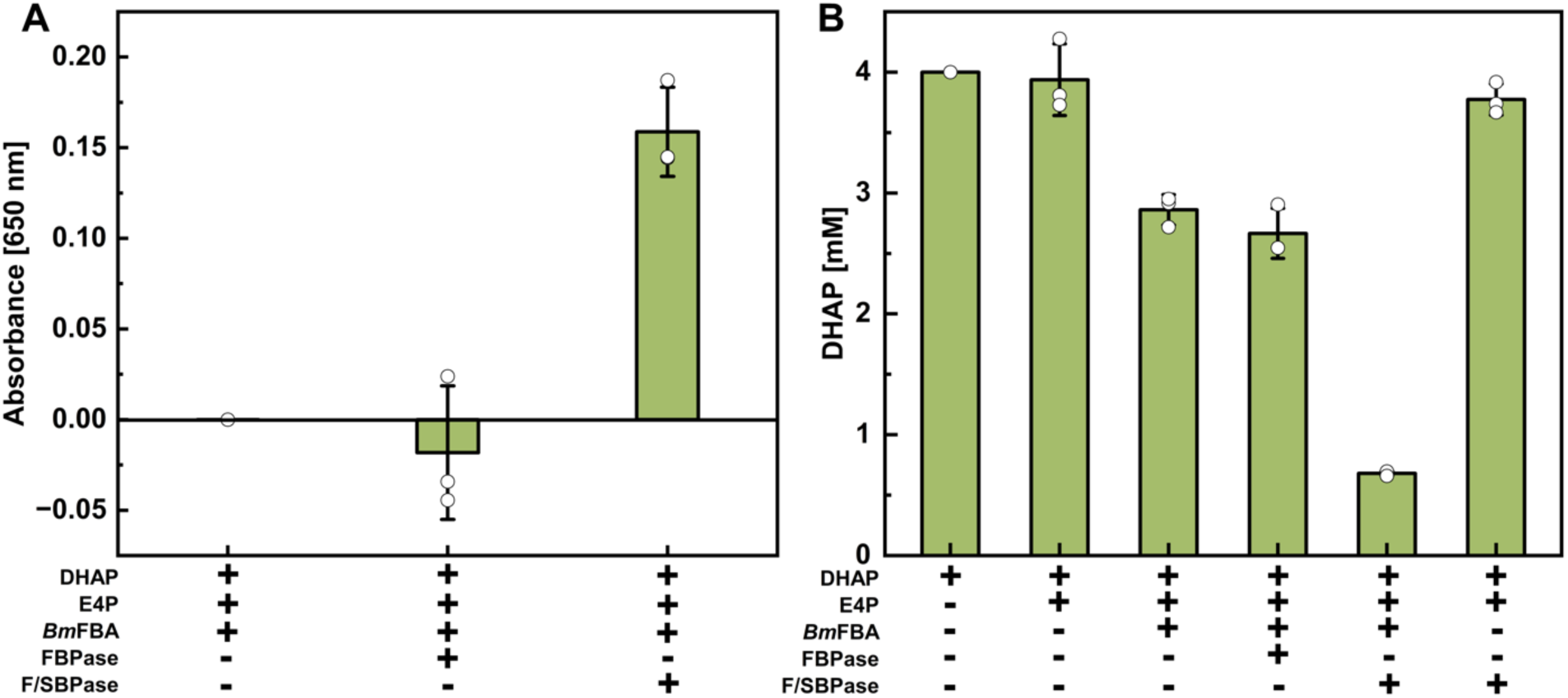
FBPase (*slr0952*) possesses no SBPase activity. **(A)** SBP was synthesized through the reversible condensation of 3 mM E4P and DHAP catalyzed by *Bm*FBA. The cleavage of SBP to S7P and PO_4_^3-^ by FBPase was assessed spectrophotometrically using a malachite green assay. The bifunctional F/SBPase (*slr2094*) served as positive control. **(B)** As an alternative to PO_4_^3-^ release, the consumption of DHAP was measured after 2 hours. DHAP levels were quantified via NADH oxidation upon the addition of 5 U G3PDH. Significant DHAP depletion, corresponding to PO_4_^3-^ and S7P formation, was observed only in the presence of both *Bm*FBA and F/SBPase, but not in the presence of FBPase. Data represent the means ± standard deviations (SD) from three technical replicates (n = 3).

### Structural comparison of *Synechocystis* FBPase and F/SBPase

Most known fructose-1,6-bisphosphatases, are metal-dependent enzymes that are classified into five distinct groups based on their amino acid sequences (Brown et al., 2009). Sequence alignments (Fig. S15) and structural comparisons using AlphaFold models of a well-characterized class I FBPase from *E. coli* (P0A993) revealed that the *Synechocystis* FBPase adopts a typical class I FBPase fold. This enzyme exhibits a high degree of sequence similarity to class I FBPases, retaining conserved residues at both the metal-binding and substrate-binding sites (Fig. 5A–C). In contrast, the conserved AMP-binding site present in *E. coli* class I FBPase (TGELT motif, highlighted in the sequence alignments and AlphaFold comparison; Fig. 5 and Fig. S15) is absent in the *Synechocystis* FBPase. Consistent with this observation, no inhibitory effect of AMP was detected at concentrations up to 10 mM (Fig. S11). Also the class I chloroplastic FBPase II (cpFBPase II) from *A. thaliana* and *Fragaria × ananassa* exhibit a similar fold to the *Synechocystis* FBPase, with the exception of an additional 20–30 amino acid sequence in the regulatory domain of the plastidial enzymes. This region, referred to as the “loop 170,” contains three cysteine residues, two of which form a disulfide bridge that is reduced by thioredoxin f during activation (Chiadmi et. al., 1999; Figs. 5 D-F and S8). Consistent with these structural differences, no regulatory effect of DTT was observed on the *Synechocystis* FBPase (Fig. S11).

**Figure 5:**
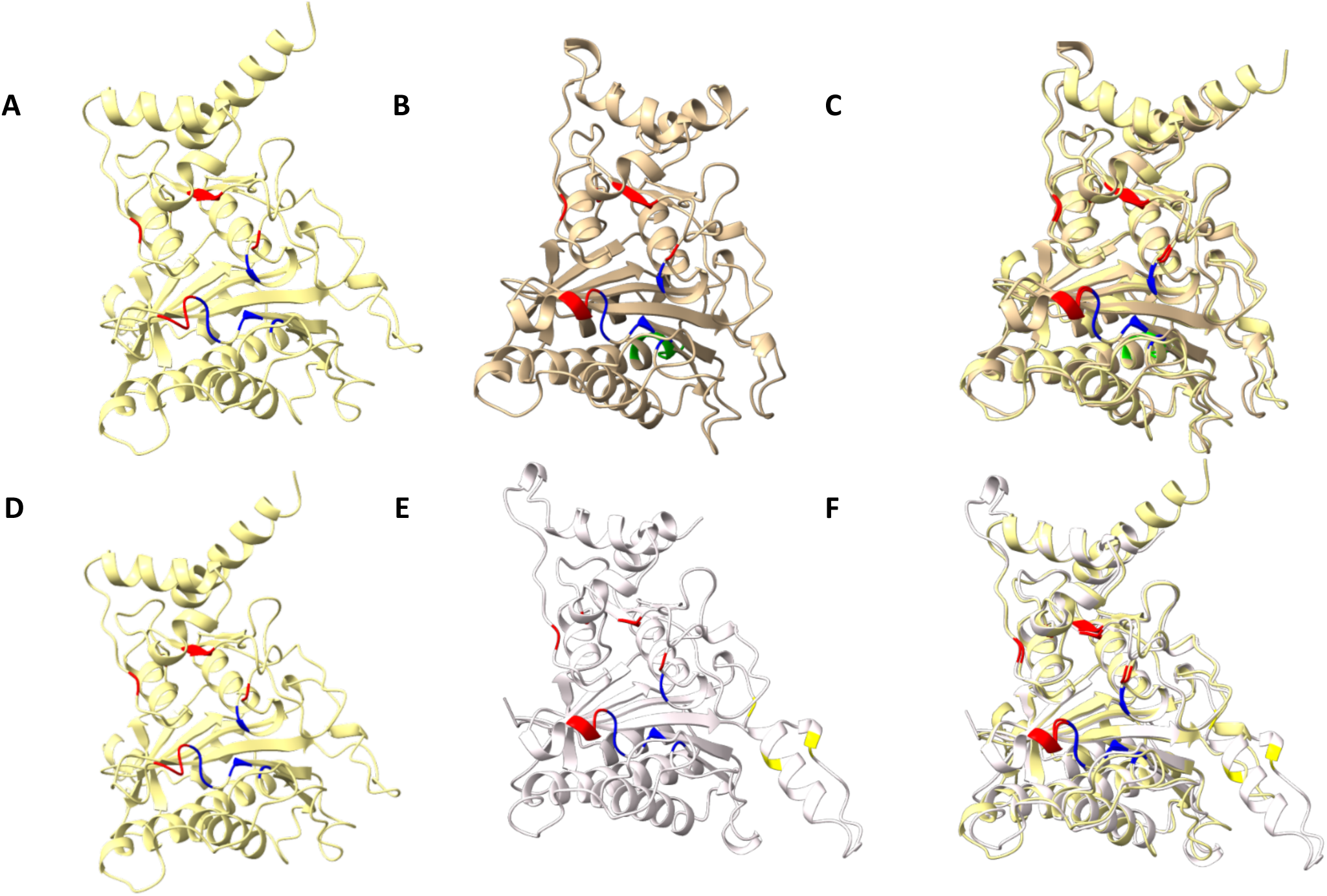
Structural comparison of the FBPase with class I FBPases from *E. coli* and *A. thaliana*. A ribbon representation (AlphaFold model) is shown for the monomer of *Synechocystis* class I FBPase (*slr0952*, yellow) **(A)** in comparison with the class I FBPase from *E. coli* (P0A993, gold, Hines et. al., 2007) **(B)**, as well as their superimposition **(C).** The AlphaFold model from *Synechocystis* class I FBPase (yellow) **(D)**, is compared to *Arabidopsis thaliana (A. thaliana*) class I cpFBPase II (P25851, grey) **(E)**, and their superimposition is shown **(F)**. Substrate, metal and AMP binding sites are highlighted in red, blue, and green, respectively. Regulatory cysteines are marked in yellow. The figures were created using AlphaFold server 3 and Chimera X.

The *Synechocystis* F/SBPase isoenzyme adopts a similar fold to that of the *E. coli* class II FBPase (Fig. S16 A–C) and is characterized by a distinct regulatory domain. This regulatory domain contains AMP-binding sites and two cysteine residues, which are involved in thioredoxin-mediated regulation, as previously described (Feng et al., 2014). Interestingly, these regulatory cysteine residues are absent in the class II FBPases of non-photosynthetic bacteria, such as *E. coli* (Brown et al., 2009).

### Phylogenetic relation of *Synechocystis* F/SBPase and FBPase to plant enzymes

We performed phylogenetic analyses to investigate the relation of *Synechocystis* F/SBPase and FBPase to enzymes from other organisms, in particular to plant cytoplasmic FBPase (cyFBPase), the two chloroplastic FBPases (cpFBPase, cpFBPase II), and the chloroplastic SBPase (cpSBPase). Sequences from green algae and proteobacteria were also included. The *Synechocystis* FBPase clusters together with FBPases from other cyanobacteria and in close proximity to proteobacterial FBPases as well as all plant enzymes (cyFBPase, cpFBPase I, cpFBPase II, and cpSBPase), which each form a distinct cluster (Fig. 6). In contrast, *Synechocystis* F/SBPase is more distantly related to the plant enzymes. It clusters with bifunctional enzymes from other cyanobacteria near the outgroup and SBPases from green algae. Overall, the tree topology indicates that (cyano)bacterial monofunctional FBPases, rather than bifunctional F/SBPases, are the origin of all plant FBPases and SBPases.

**Figure 6:**
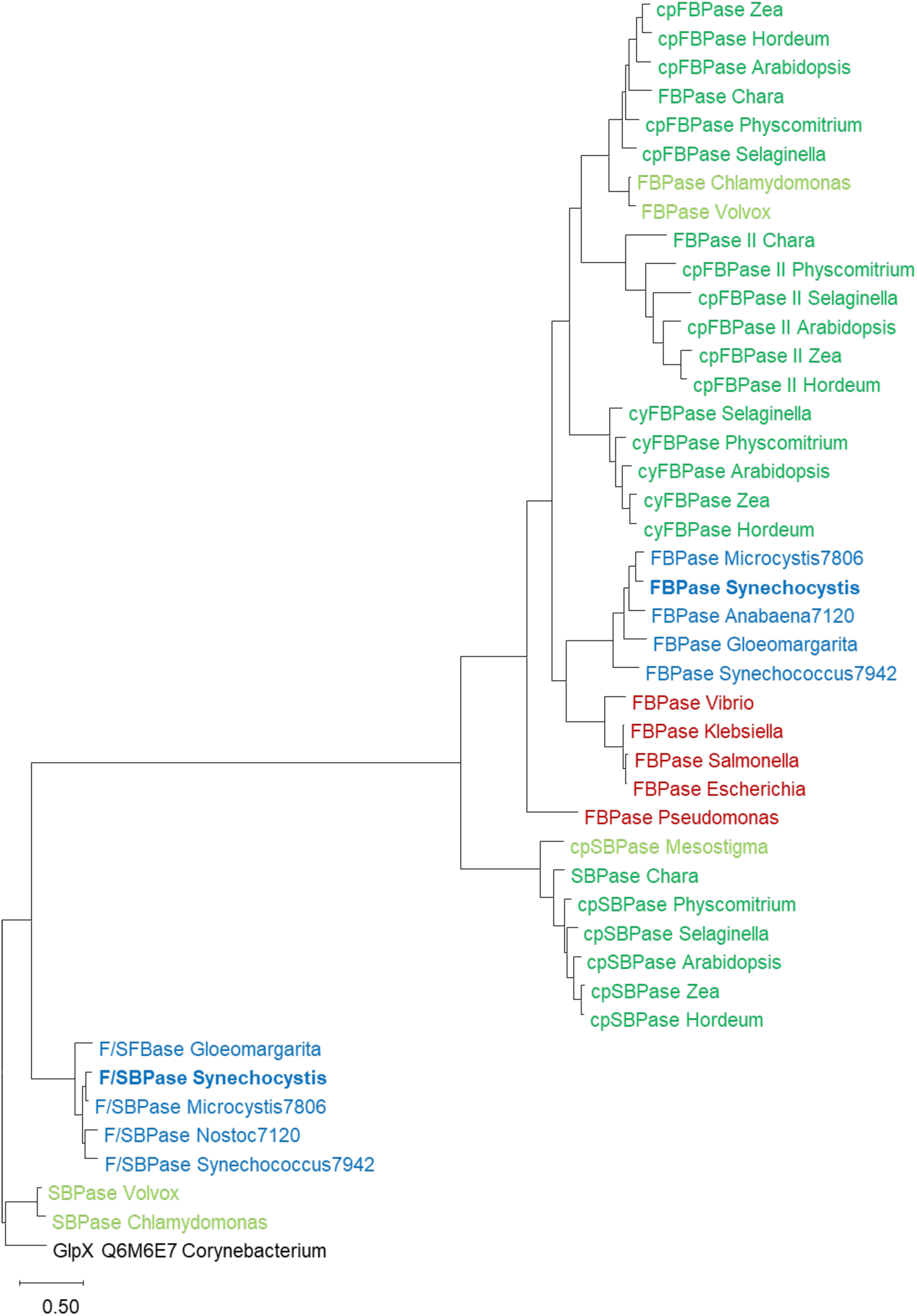
Phylogenetic tree of FBPases and SBPases from photoautotrophic and heterotrophic organisms. Enzymes from cyanobacteria are colored in blue (*Synechocystis* in bold), from proteobacteria in red, from chlorophyta (green algae) in light green, and from streptophyta (plants) in dark green. As outgroup, the multifunctional sugar phosphatase (GlpX) from *Corynebacterium* was used. The tree was generated in MEGA with a maximum-likelihood analysis of aligned and trimmed sequences. Accession numbers of all sequences used in the phylogenetic cluster analysis can be found in the Supplementary Table S2.

### *In vivo* FBPase activity in deletion mutants

To analyze the *in vivo* function of FBPase in a cellular context, a collection of deletion mutants was created. In contrast to other groups (García-Cañas et al., 2022; Yan & Xu, 2008), our attempts to segregate the *slr2094* deletion (Δ*f/sbpase*) were not successful, similar to earlier studies which reported the deletion of F/SBPase to be lethal (Tamoi et al., 1999). However, for *slr0952* (Δ*fbpase*) complete deletion was successful (Figs. S1-3, Supplemental Text S1). In the following, a merodiploid *f*/*sbpase*/Δ*f*/*sbpase* and a fully segregated Δ*fbpase* mutant were characterized.

First, we analyzed the *in vivo* contribution of the two enzymes on total FBPase activity in crude cell extracts by performing an enzyme activity assay with FBP as substrate. As shown above, bifunctional F/SBPase in contrast to FBPase has to be activated by reducing conditions. Therefore, cells were disrupted under either reducing or under oxidizing conditions, enabling us to distinguish the activities of the two enzymes in the different strains.

For all strains, FBPase activity under reducing conditions was much higher than under oxidizing conditions (p < 0.05 for each strain) (Fig. 7). This can be explained by the specific activation of F/SBPase by reduction, which is expressed at a much higher level than FBPase (Saha et al., 2016). Under reducing conditions FBPase activity was similar in WT, *f*/*sbpase*/Δ*f*/*sbpase* and Δ*fbpase*. The fact that we could not detect a difference in total cellular FBPase activity between the WT and the Δ*fbpase* mutant indicates that the activity of FBPase is negligible compared to that of F/SBPase under those conditions. The WT-like FBPase activity of the merodiploid *f*/*sbpase*/Δ*f*/*sbpase* mutant can likely be attributed to an upregulation of the expression of the remaining copies of *f/sbpase*. An upregulation of *fbpase* expression cannot explain this observation, because also under oxidizing conditions the *f*/*sbpase*/Δ*f*/*sbpase* mutant showed the same activity as the WT and FBPase is not activated by reduction. In previous studies FBPase activity was virtually lacking in a fully segregated *f/sbpase* deletion mutant, which led to the conclusion that F/SBPase might be the only active FBPase *in vivo* (Yan & Xu, 2008). Interestingly, we observed that deletion of *fbpase* resulted in a significant reduction of FBPase activity by around two thirds under oxidizing conditions (p < 0.001, Fig. 7). This suggests that, unlike previously assumed, the monofunctional FBPase is indeed active under oxidizing conditions *in vivo*, albeit at low rates. The remaining activity in Δ*fbpase* under oxidizing conditions can likely be attributed to residual activity of F/SBPase due to incomplete inactivation.

**Figure 7:**
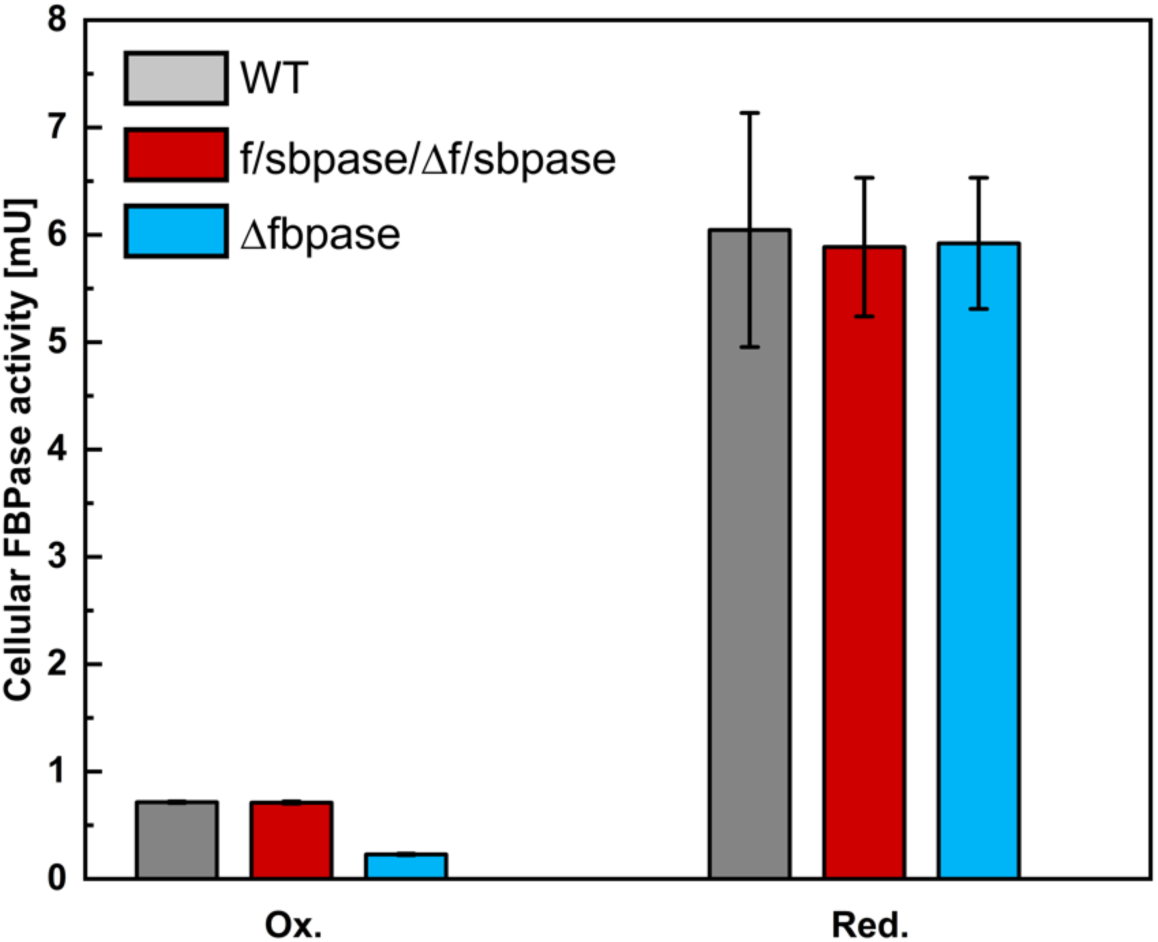
Cellular FBPase activity in deletion mutants. FBPase activity was measured in crude cell extracts of WT, the merodiploid *f*/*sbpase*/Δ*f*/*sbpase* mutant and Δ*fbpase* that were obtained under either oxidizing (Ox.) or reducing (Red.) conditions and normalized to the same phycobilisome concentration. Mean, maximum and minimum values of 2 (WT) or 3 (*f*/*sbpase*/Δ*f*/*sbpase* and Δ*fbpase*) biological replicates are depicted.

### Growth of *f/sbpase, fbpase* and *talB* deletion mutants

To further investigate the physiological role of the two F(/S)BPases, we investigated the growth of the *fbpase* and merodiploid *f/sbpase* deletion mutants under different conditions. Additionally, we included deletion mutants of *talB* (encoding the OPP pathway-specific transaldolase) to better understand its relation to FBPase and F/SBPase, because the enzyme TalB can theoretically revert the reaction of SBPase (see Fig. 1).

Growth of the merodiploid *f*/*sbpase*/Δ*f*/*sbpase* mutant was not affected under any of the light conditions we tested, which is well in line with the interpretation that this mutant upregulated F/SBPase activity of the bifunctional enzyme to WT levels (compare Figs. 7, 8A-D). Similarly, the Δ*fbpase* mutant grew like the WT under photoautotrophic and photomixotrophic conditions in constant light (Fig. 8A, C). These results are consistent with previous studies, which showed that the absence of FBPase affected neither photoautotrophic, photoheterotrophic nor photomixotrophic growth, whereas F/SBPase was essential under photoautotrophic conditions (García-Cañas et al., 2022). Furthermore, in diurnal day/night cycles photoautotrophic and photomixotrophic growth was not affected in Δ*fbpase* (Fig. 8B, D). However, under heterotrophic conditions (darkness with glucose supplied), we observed differences. The growth of *f*/*sbpase*/Δ*f*/*sbpase* was faster than that of WT, whereas surprisingly that of Δ*fbpase* was diminished (Fig. 8E). These results indicate that FBPase, in contrast to the role of F/SBPase in the CBB cycle during the day, plays a role in the dark rather than in the light. This fits well with our biochemical data, which show that FBPase retains its activity under oxidizing conditions, whereas F/SBPase is activated under reducing conditions and inhibited by AMP (low energy charge; Figs. 7, S4, S5). The likely role of FBPase in dark phases is further supported by the fact that its expression is upregulated during the night in contrast to F/SBPase (Saha et al., 2016).

**Figure 8:**
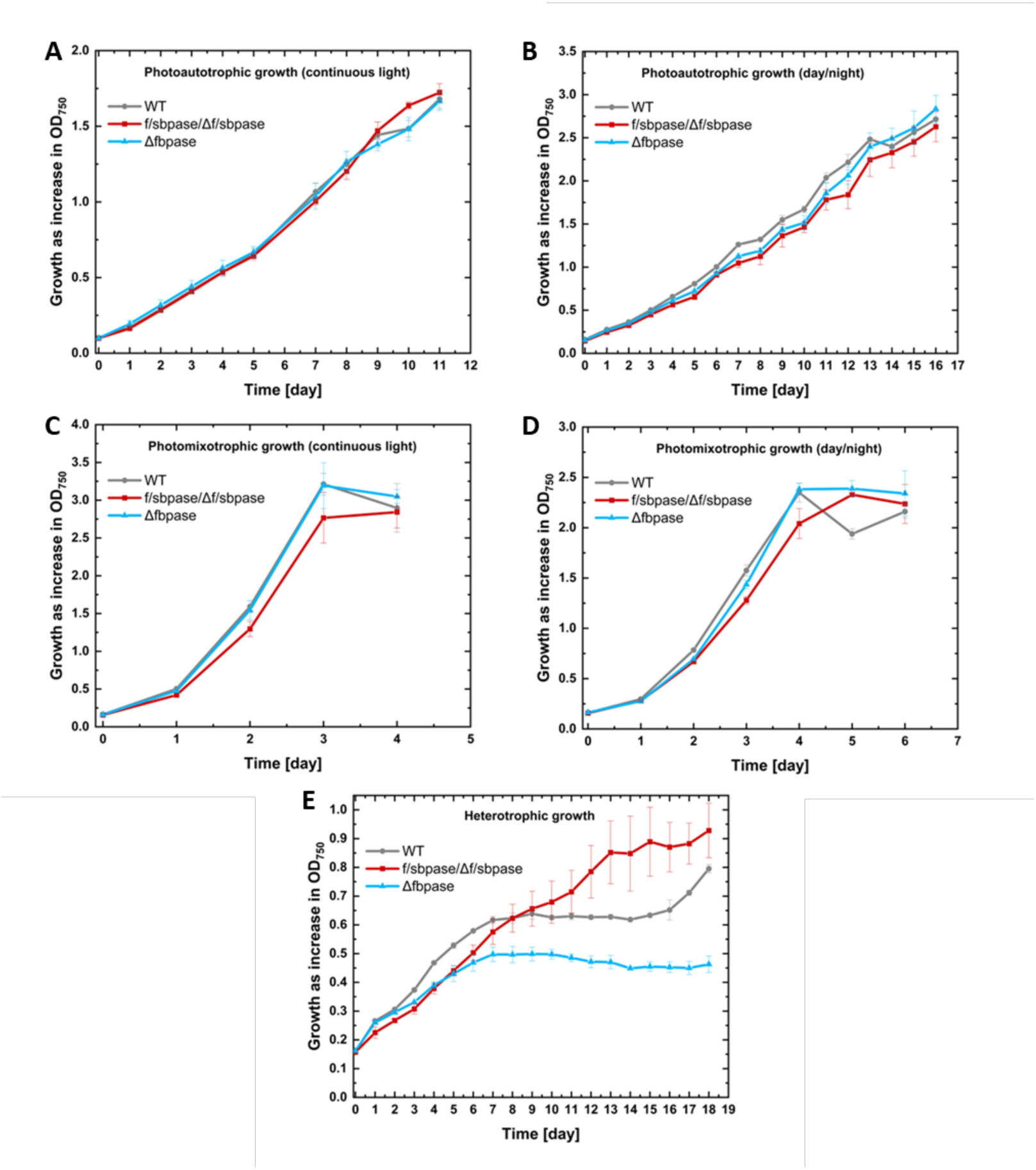
Growth of Δ*@pase* and merodiploid *f/sbpase/*Δ*f/sbpase* deletion mutants compared to WT. Photoautotrophic and photomixotrophic growth in continuous light **(A, C)** and day/night cycles **(B, D)**, and in light activated heterotrophic growth **(E)** were measured. Mean, maximum and minimum of 3 biological replicates are depicted. Each experiment was conducted at least 3 times and the results of one representative repetition are shown.

Deletion of *talB* did not affect photoautotrophic or photomixotrophic growth in continuous light, or phototrophic growth in day/night cycles, neither in the single mutant nor in the double mutants in combination with *f*/*sbpase*/Δ*f*/*sbpase* or Δ*fbpase* (Fig. S17A-C). However, growth of mutants with deleted *talB* was diminished under photomixotrophic conditions in day/night cycles (Fig. S17D) and completely abolished under heterotrophic conditions (Fig. S17E). Additional deletion of *f*/*sbpase*/Δ*f*/*sbpase* or Δ*fbpase* did not suppress growth further. On the contrary, the Δ*talB*Δ*fbpase* seemed to grow slightly better than the Δ*talB* single mutant in photomixotrophic day/night cycles. The observation that deletion of *talB* only impacts growth in conditions with dark phases indicates that the enzyme is confined to its role in the OPP pathway in darkness and is rather not involved in the CBB cycle in light in *Synechocystis*.

### Growth of *pfk* deletion mutants

To clarify the relevance of the two isoenzymes of PFK and their relation to FBPase, we conducted growth experiments with WT, the double (Δ*p[*) and single deletion mutants (Δ*p[-A1,* Δ*p[-A2*). Under photoautotrophic and photomixotrophic conditions, Δ*p[* grew like the WT but was impaired in growth under heterotrophic conditions (Fig. 9), consistent with previous studies (Makowka et al., 2020). As expected, also both single deletion mutants grew like the WT under photoautotrophic and photomixotrophic conditions (Fig. 9A, B). Under photoheterotrophic conditions the double and single *p[* deletion mutants grew again similar as the WT, however, Δ*p[-A2* reached reproducibly lower optical densities (Fig. 9C). Under heterotrophic conditions, Δ*p[-A2* growth was more strongly affected than that of Δ*p[-A1* (Fig. 9D). Interestingly, the growth of the double deletion mutant lay between that of the two single deletion mutants.

**Figure 9:**
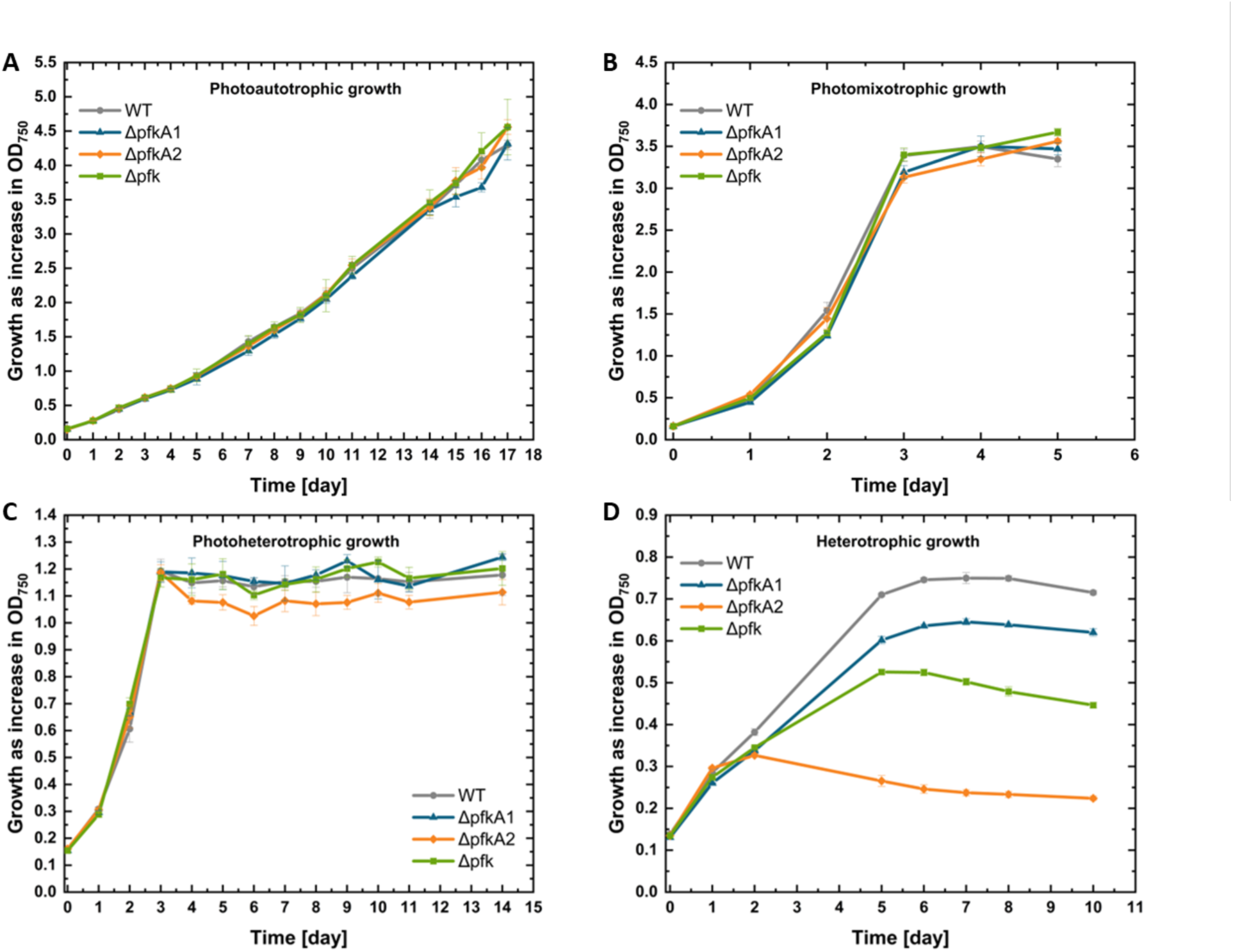
Growth of the *pfi* single and double deletion mutants compared to WT under different conditions. Photoautotrophic **(A)**, photomixotrophic **(B)**, and photoheterotrophic **(C)** growth in continuous light, and light activated heterotrophic growth **(D)** were measured. The double deletion mutant Δ*p[-A1*Δ*p[-A2* is abbreviated as Δ*p[*. Mean, maximum and minimum of 3 biological replicates are depicted. Each experiment was conducted at least 3 times and the results of one representative repetition are shown.

Apparently both PFK isoenzymes are active and required for optimal growth in the dark with external glucose. They do not seem to be able to (fully) compensate the absence of each other, even though PFK-A2 seems to be more important.

## Discussion

In summary, our results provide the following comprehensive picture of the switch of the cyanobacterial carbohydrate metabolism between light and darkness, which is shown in Figure 10.

**Figure 10:**
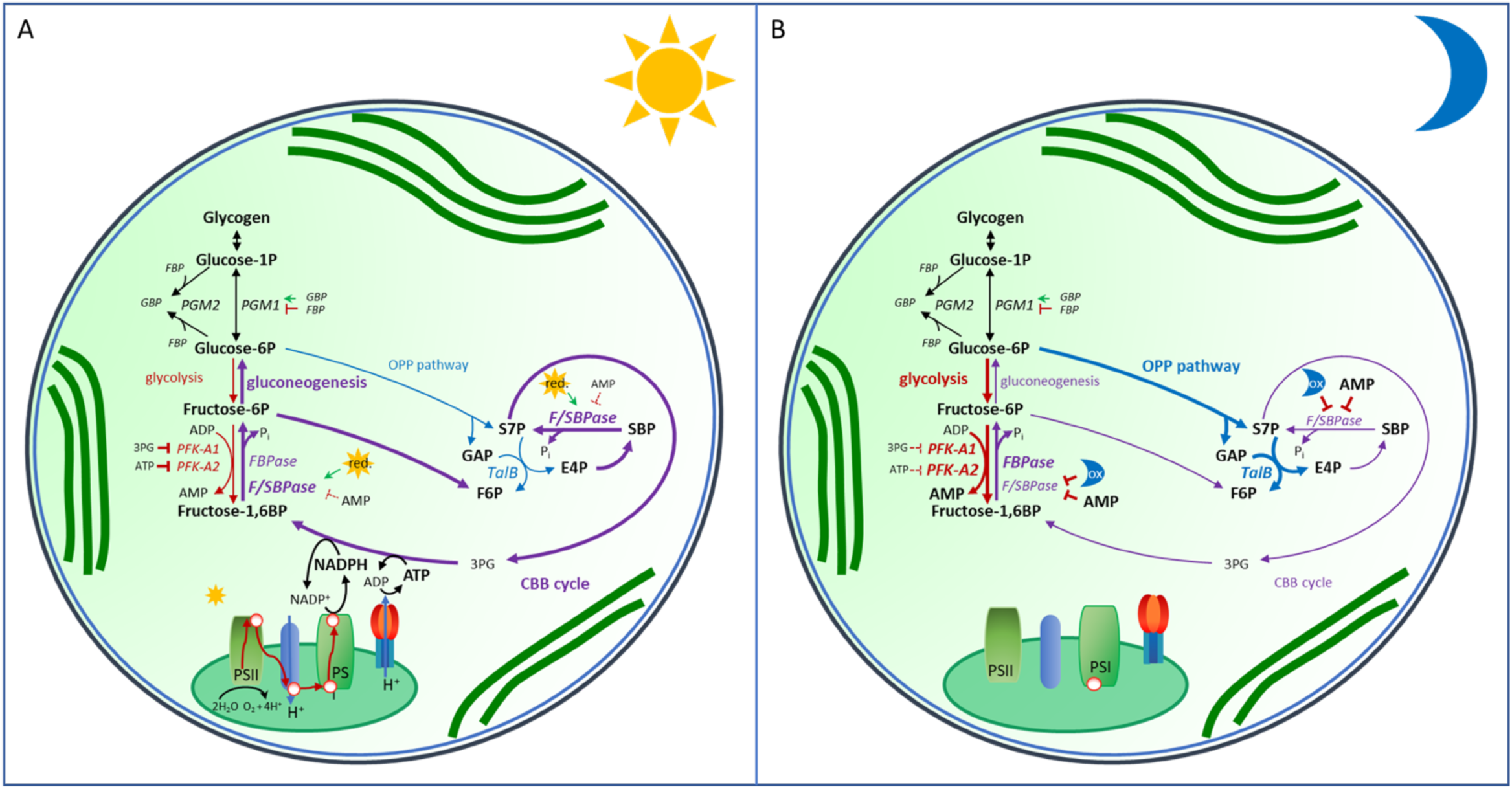
Overview of the cyanobacterial carbohydrate metabolism in light and darkness. **(A)** In light the photosynthetic light reaction, as well as the anabolic reactions of CBB cycle and gluconeogenesis are active, the bifunctional F/SBPase catalyzing two key reactions. **(B)** Under heterotrophic conditions in darkness the carbon flow is reversed to the catabolic direction via the OPP pathway (depending on TalB) and glycolysis (supported by both PFKs), although FBPase is active in parallel. The activation of F/SBPase in the light and PFKs and FBPase in darkness is achieved by their different regulation, permitting the modulation of fluxes to environmental conditions and metabolic needs of the cell. (FBP = fructose-1,6-bisphosphate; F6P = fructose-6-phosphate; GBP = glucose-1,6-bisphosphate; 3PG = 3-phospho-glycerate; E4P = erythrose-4-phosphate; SBP = seduheptulose-1,7-bisphosphate; GAP = glyceraldehyde-3-phosphate; S7P = seduheptulose-7-phosphate; PGM = phosphoglucomutase; TalB = transaldolase; FBPase = fructose-1,6-bisphosphatase; F/SBPase = fructose-1,6-biphosphatase/sedoheptulose-1,7-biphosphatase; PFK = phosphofructokinase)

Under photoautotrophic conditions, the light reaction of photosynthesis provides ATP and NAPDH to fuel the CBB cycle. F/SBPase is activated by reduced thioredoxin. AMP levels are probably below F/SBPase inhibiting concentrations in light. We showed that FBPase does not possess SBPase activity and its deletion does not affect photoautotrophic growth. Therefore, F/SBPase is most likely the sole enzyme responsible for dephosphorylating not only SBP, but also FBP to drive the CBB cycle in direction of ribulose-1,5-bisphosphate synthesis and gluconeogenesis in the light. This interpretation is furthermore supported by a previous study in which only the simultaneous expression of plant cpSBPase and cpFBPase restored photoautotrophic growth in the fully segregated Δ*f/sbpase* mutant (García-Cañas et al., 2022). The cell’s own FBPase was apparently unable to replace the missing FBPase activity of the deleted *f/sbpase* in the light. Based on our measurements of FBPase activity in crude cell extracts, a likely explanation is that its activity in reducing conditions is negligible compared to that of F/SBPase. The low activity of FBPase may, among other reasons, result from its approximately 30-fold lower protein abundance compared to F/SBPase (Jackson et al., 2022).

The growth behavior of various *talB* deletion mutants verifies that this enzyme only plays an essential role under heterotrophic conditions and is confined to the OPP pathway. While some chemoautotrophic bacteria that lack F/SBPase can use transaldolase instead (Frolov et al., 2019), *Synechocystis* is apparently unable to replace F/SBPase with transaldolase. Attempts to delete *f/sbpase* either result in mutants that were no longer able to grow photoautotrophically (García-Cañas et al., 2022; Yan & Xu, 2008) or the segregation of the mutants did not succeed (Tamoi et al., 1999). TalB is therefore unable to functionally replace F/SBPase in the CBB cycle.

Similarly, both PFK isoenzymes are inactive in the light, as evidenced by the growth behavior of the deletion mutants. The relatively high level of ATP produced in the photosynthetic light reaction inhibits PFK-A2, while 3PG, the first stable product of photosynthetic CO_2_ fixation by Rubisco, inhibits PFK-A1 (Shen et al., 2024). Additionally, the concentration of their co-substrate ADP is relatively low. By utilizing ADP instead of ATP, the latter being the common co-substrate of PFKs from heterotrophic organisms, activation of PFKs is prevented under photoautotrophic conditions in cyanobacteria (Shen et al., 2024).

In the dark, oxidizing conditions prevail, so that F/SBPase is inactivated. Due to the absence of the photosynthetic light reaction the energy charge is presumably low in darkness, removing the inhibition of PFK-A2 by ATP and providing ADP as co-substrate for both PFKs. The inactivity of the CBB cycle results in lower 3PG concentrations, which removes inhibition of PFK-A1 (Shen et al., 2024). The activity of PFKs provides AMP, which further supports inhibition of F/SBPase. Thereby, the carbon metabolism is switched from an anabolic to a catabolic direction via glycolysis and OPP pathway. Flux analyses show that under heterotrophic conditions carbohydrates are primarily metabolized via the OPP pathway, while glycolysis is used to a lesser extent (Wan et al., 2017; Yang et al., 2002). This explains the strong effect of *talB* deletion on heterotrophic growth. Accordingly, the PFKs are less important, although both PFK-A1 and PFK-A2 are required for optimal heterotrophic growth. Possessing two isoenzymes might have the advantage of providing greater flexibility and enabling a finer regulation of metabolism (Jablonsky et al., 2014).

Even though catabolic pathways are operating in darkness, the anabolic FBPase is apparently nevertheless active as well, as its deletion leads to a reduction of growth. This is further supported by the fact that it is not inactivated by oxidation and its expression is upregulated in darkness, compared to light (Saha et al., 2016). The question arises which function the FBPase could fulfill. As an antagonistic enzyme couple FBPase and the ADP-dependent PFK-As catalyze exergonic irreversible biochemical reactions between two substrates (F6P and FBP) in opposite directions and their simultaneous operation results in an energy dissipating futile cycle. Such substrate cycles between PFKs and FBPases are well known from eukaryotes for thermogenesis (Sharma et al., 2024; Staples et al., 2004). Another assumed function of substrate cycles is that they permit a better fine-tuning of metabolic fluxes by enhancing the sensitivity of enzymes to changes in effector concentrations (Newsholme et al., 1984). In particular, switching between zero to moderate flow rates can be achieved more easily in the presence of regulated forward and back reactions. The *Synechocystis* FBPase could therefore have the task of fine-tuning the flow through the EMP pathway under heterotrophic conditions by catalyzing its back reaction. Additionally, adjusting the concentration of its substrate FBP could play a regulatory role, for instance in glycogen metabolism. FBP has been shown to directly inhibit phosphoglucomutase 1 (PGM1), which catalyzes the interconversion of glucose-1-phosphate (G1P) and G6P in glycogen metabolism. A second phosphoglucomutase 2 (PGM2) however, uses FBP as a phosphodonor to produce GBP with either G1P or G6P (Neumann et al., 2022). GBP in turn is an essential activator of PGM1. FBP thus has counteracting effects on the activity of PGM1. On the one hand it acts as a direct inhibitor of PGM1, on the other hand it is used for the formation of GBP by PGM2, which activates PGM1 (Neumann et al., 2022). We found that GBP cannot be metabolized by FBPase. However, FBPase could indirectly control GBP levels by adapting the concentrations of FBP, the substrate of PGM2. In collaboration with the PFKs, FBPase could thus serve the purpose of regulating the flow of glycogen into glycolysis via PGM1 under heterotrophic conditions. Further studies are required to substantiate this hypothesis.

Phylogenetic analyses revealed that *Synechocystis* FBPase is more closely related to all plant FBPases and SBPase than *Synechocystis* F/SBPase. The redox regulation in *Synechocystis* F/SBPase and plant cpFBPase might thus be the result of convergent evolution and an adaptation to a photoautotrophic lifestyle. The second chloroplastic FBPase (cpFBPase II) is redox-insensitive like *Synechocystis* FBPase (Li et al., 2020; Serrato et al., 2009). Therefore, it would be interesting to test if they might fulfil a similar function in dark metabolism.

In conclusion, F/SBPase and FBPase fulfill distinct functions in the cyanobacterium *Synechocystis*. While F/SBPase is highly abundant and regulated by different biochemical effectors (AMP and redox charge) ensuring its activation in the light and inactivation in darkness, FBPase is less abundant and is not regulated by effectors but rather on a transcriptional level, elevating its level under dark conditions. Hence, F/SBPase catalyzes reactions in CBB cycle and gluconeogenesis in the light, whereas FBPase plays a role under heterotrophic conditions in darkness, probably by forming a regulatory futile substrate cycle with the PFKs. The antagonistic enzyme pair of FBPase and PFKs might fulfil the function to fine-tune the flux through the EMP pathway and/or to control glycogen metabolism via FBP levels and PGM1. The classical control point in the EMP pathway, which is known to be mediated by the antagonistic enzyme pair PFK and FBPase in heterotrophic bacteria and eukaryotes, is therefore also present in *Synechocystis*. However, adaptation to its dual heterotrophic and photoautotrophic lifestyle demands a more intricate regulatory mechanism, involving two ADP-dependent PFK-A isoenzymes, one FBPase, and a bifunctional F/SBPase. We furthermore show that transaldolase is confined to the OPP pathway in *Synechocystis*, most likely not being involved in the CBB cycle, and that deletion of *p[-A2* affects heterotrophic growth to a larger extent than deletion of *p[-A1*. The complex network of differently expressed and regulated isoenzymes at the control points for anabolic and catabolic carbon metabolism enables cyanobacteria to switch the direction of carbon flow and to precisely regulate its extent, based on light conditions.

## Abbreviations

2PG: 2-phosphoglycolate
3PG: 3-phosphoglycerate
CBB: cycle Calvin-Benson-Bassham cycle
DCMU: 3-(3,4-dichlorophenyl)-1,1-dimethylurea
DHAP: dihydroxyacetone phosphate
E4P: erythrose-4-phosphate
EMP: pathway Embden-Meyerhoff-Parnass pathway
F1P: fructose-1-phosphate
F6P: fructose-6-phosphate
FBA: fructose-1,6-bisphosphate aldolase
FBP: fructose-1,6-bisphosphate
FBPase: fructose-1,6-bisphosphatase
G3PDH: glycerol-3-phosphate dehydrogenase
G1P: glucose-1-phosphate
G6P: glucose-6-phosphate
G6PDH: glucose-6-phosphate dehydrogenase
GAP: glyceraldehyde-3-phosphate
GBP: glucose-1,6-bisphosphate
IMAC: metal ion affinity chromatography
NOPP: pathway non-oxidative pentose phosphate pathway
OPP: pathway oxidative pentose phosphate pathway
PEP: phosphoenolpyruvate
PFK: phosphofructokinase
PGI: phosphoglucose isomerase
PGM: phosphoglucomutase
PHB: polyhydroxybutyrate
Ru5P: ribulose-5-phosphate
RPP: pathway reductive pentose phosphate pathway
S7P: seduheptulose-7-phosphate
SBP: seduheptulose-1,7-bisphosphate
SBPase: sedoheptulose-1,7-biphosphatase
SEC: size exclusion chromatography
TalB: transaldolase
WT: wildtype

## Supplementary data

The following supplementary data are available at JXB online.

Table S1: Primer list

Table S2: Accession numbers of protein sequences used for phylogenetic analyses

Table S3: Mutant list

Figure S1: Segregation for *f/sbpase* in the single and double *f(/s)bpase* deletion mutants

Figure S2: Segregation for *f/sbpase* in different cultures of the *f/sbpase* single deletion mutant and of the *ΔtalB f/sbpaseΔf/sbpase* double deletion mutant

Figure S3: Segregation for *fbpase* in the single and double deletion mutants

Figure S4: Segregation of the *talB* single deletion mutant and the *talB f/(s)bpase* double deletion mutants

Figure S5: Segregation of the *p[* single and double deletion mutants

Figure S6: Test of redox conditions in the FBPase activity assay with cell extracts

Figure S7: FBpase activity in cell extracts under different redox conditions

Figure S8: Purification of the recombinant Synechocystis FBPase (*slr0952*)

Figure S9: Native molecular mass of FBPase

Figure S10: Metal dependency and optimal concentration for FBPase activity

Figure S11: FBPase activity in the presence of ATP, AMP and DTT

Figure S12: Enzymatic activity and regulatory properties of F/SBPase (*slr2094*)

Figure S13: NMR spectra of FBPase activity by FBPase and F/SBPase

Figure S14: Glucose-1,6-bisphosphate interaction with FBPase and F/SBPase

Figure S15: Sequence alignment of FBPase with class I FBPase from *E. coli*, cpFBPase I and cyFBPase I

Figure S16: AlphaFold comparisons of F/SBPase with *E. coli* class 2 FBPase and *Synechocystis* FBPase

Figure S17: Growth of the *talB* single and double deletion mutants

Supplemental Text S1: Mutant segregation

Supplemental Text S2: Verification of the redox-regulation of F/SBPase observed in cell extracts

## Acknowledgements

We would like to express our sincere gratitude to Prof. Volker F. Wendisch (Universität Bielefeld) for providing the pET16b_bmmga3_16125 plasmid and Rudolf Walter for his valuable experimental contributions to this research project.

## Author contributions

FC, RO, LS, MG, MB performed the experiments, FC, RO, LS, CB, JLS, KF, MH, BS, KG analyzed data, FC and KG wrote the original draft, RO, BS, MH, wrote, reviewed and edited, KF, MH, BS, KG conceptualized the study, BS and KG supervised, KF, MH, BS, KG funding acquisition.

## Conflict of interest

The authors declare that they have no conflicts of interest with the contents of this article.

## Funding

We acknowledge funding by the German Research Foundation, Bonn (Deutsche Forschungsgemeinschaft, DFG) for the Research Unit SCyCode (FOR2816) (grant SI 642/ 14–1 and SI 642/14–2 to B. S., HA2002/23–2 to M. H., GU 1522/5–1 to K. G., and FO195/16–2 to K. F.), as well as financial assistance from the DSI/NRF in South Africa (grant NRF-SARCHI-82813 to J. L. S.).

## Notes

### Competing Interest Statement

The authors have declared no competing interest.

